# The prion-like protein Doppel: A soluble biomarker steering ovarian cancer’s peritoneal to circulatory dissemination

**DOI:** 10.1101/2024.07.26.605386

**Authors:** Zulfikar Azam, Xiaojun Zhang, Riajul Wahab, Md Mahedi Hasan, Bowon Kang, Md Mynul Hassan, Mazharul Karim, Jeong Uk Choi, Muhit Rana, Jiya-Ying Zhang, Sourav Roy, Youngro Byun, In-San Kim, Jae Yun Song, Farzana Alam, Eugene P. Toy, Sireesha Y. Reddy, Taslim A. Al-Hilal

## Abstract

Detecting ovarian cancer (OC) early using existing biomarkers, e.g., cancer antigen 125 (CA125), is challenging due to its ubiquitous expression in many tissues. Doppel, a prion-like protein, expresses in male reproductive organ but absent in female reproductive systems and healthy tissues, but plays an important role in neoangiogenesis. Here, we have shown two platforms, soluble Doppel in sera/ascites and Doppel expressed circulating tumor cells (^Dpl+^CTC) in the whole blood, to detect subsets of epithelial OC (EOC). Increased level of Doppel in the sera of OC patients, in three different cohorts, confirm Doppel as OC specific biomarker. Serum Doppel level distinguishes EOC subtypes and early stages HGSOCs from non-cancerous conditions with high sensitivity and specificity. Stratifying the EOCs based on Doppel level, we categorized them into Doppel-high (Dpl^hi^) and Doppel-low (Dpl^low^) groups. Using ascites-derived organoids and single cell sequencing of whole ascites of Dpl^hi^ and Dpl^low^ patients, we identify that Doppel induces epithelial-mesenchymal transition (EMT) and creates an immunosuppressive microenvironment, respectively. Doppel levels in the sera/ascites correlate with the changes of ^Dpl+^CTC number in whole blood, highlighting the association of Doppel-induced EMT with CTC dissemination in circulation. Thus, Doppel-based detection of EOC subtypes could be a promising platform as clinical biomarker and link Doppel-axis with OC dissemination.

## Introduction

The latent symptoms and subtle onset of ovarian cancer (OC) pose a significant challenge in identifying the disease at its earlier stages^[1]^. Frequent metastasis and failure to detect earlier exacerbate the OC progression and contribute greatly for OC patient’s overall survival, which stands at less than 20%, 5 years survival from disease diagnosis for stage III and IV OC patients^[2]^. The high morbidity and mortality of OC patients underscores the importance of identifying new biomarkers which can effectively detect and stratify the course of OC cancer fate. Beyond the detection of OC cancer early, diagnostic methods are also needed to track disease progression, treatment response and treatment resistance. The cancer antigen 125 (CA-125) and human epididymis protein 4 (HE4), two FDA approved serum glycoprotein biomarkers, are widely used to detect patients with transvaginal ultrasound (TVUS). The utility of CA-125 as reliable biomarker for OC diagnosis is less certain than expected. As a glycoprotein CA-125 is widely expressed in normal tissue types along with non-cancerous conditions in women, making it less specific for OC diagnosis ^[3]^. The sensitivity of CA-125 is not sufficient to detect epithelial ovarian cancer (EOC) with one in five found significantly lower level of CA-125 ^[4]^. HE4’s ability to distinguish EOC from benign masses is superior to CA-125. But the level of HE4 is increased by multiple non-ovarian influences and various malignancies, predominantly those originating from the reproductive system but also encompassing respiratory cancers ^[5]^ ^[6]^. Thus, the discovery of potential selective biomarker that is absent in the sera or reproductive organs of healthy female but present in ovarian cancer patients are immensely needed.

Doppel, encoded by the *PRND* gene, is a 179-amino-acid polypeptide with domains similar to those of cellular prions (PrP), sharing 25% structural homology with PrP ^[7]^. Based on amino acid pairing, human Doppel has 79% similarity with mouse Doppel ^[7]^. Doppel was first identified in 1999 and later discovered to be expressed in human adult testis and therefore is believed to play a role in male fertility ^[8]^ ^[9]^. Recent quantitative transcriptomic analysis (RNA-seq) of the human genome confirmed the testis-specific expression of Doppel ^[10]^. Although the brain endothelium of newborn mice expresses Doppel transiently, the endothelium of adult mouse brain expresses no Doppel ^[11]^. Doppel knock-out mice, created from full immunogenic mice, exhibit no developmental defects except sterility in males, suggesting that Doppel may have no role in the development and physiology of female reproductive systems^[12]^. We and others reported that Doppel is a highly specific tumor endothelium marker ^[13]^ ^[14]^. We showed that both human and mouse tumors express Doppel in their vasculatures, inhibition of Doppel depletes cell-surface VEGFR2, and thus abrogates VEGF binding with the receptor ^[14]^.

In this study, we seize the opportunity of Doppel that is absent in female reproductive organ as well as in any healthy tissues other than testis, to establish Doppel as an OC diagnostic biomarker by utilizing clinical specimens (serum and ascites). We elaborated our findings to correlate Doppel’s role in OC dissemination into ascites and blood by identifying circulating tumor cells (CTCs), and its relationship with immune modulator by utilizing the power of patient-derived organoids and single cell RNA sequencing.

## Results

### Doppel expression is linked to poor prognosis and survival in human epithelial ovarian cancers

We sought to determine the expression level of Doppel in OC patients, using public RNA sequencing data of GTEX normal ovary tissue (n=88) and TCGA ovarian primary serous adenocarcinoma (n=419), and result strongly indicated the association of Doppel in OC pathobiology **(Fig 1A).** In line with expression differences, the survival differences between *PRND*-high and *PRND*-low serous OC groups showed significant differences both overall and progression free survival **(Fig 1B and 1C)** clearly indicating Doppel’s usefulness as survival indicator of OC. We next aimed to benchmark Doppel in OC tissues using tissue microarray which included 54 cases of HGSOC, 8 normal adjacent tissues (NAT), and 2 ovary tissues. Despite some variability, 83% (45 of 54 samples) of HGSOC of ovarian cancers showed high Doppel expression (**Fig. 1D**), whereas none of the NAT and normal ovary tissues were found to be positive for Doppel **(Fig. 1E)**. Doppel was predominantly found in the tumor vasculatures of HGSOC, but not in the NAT (**Fig. 1F**). Normal endothelial cells of tumor-free regions and tumor endothelial cells were isolated from fresh tissues immediately after surgical debulking of ovarian tumors, according to our laboratory standard protocol ^[14]^. Doppel protein and mRNA level is found high in TECs of all three HGSOC patients (**Fig. 1G and 1H**). This finding aligned with our previous findings ^[14]^ where we demonstrated the expression of Doppel mainly localized in tumor associated endothelial cells in primary tissues. Together, these findings indicate Doppel is prognostic of poor survival and tissue expression of Doppel is abundant in OC.

**Figure 1:**
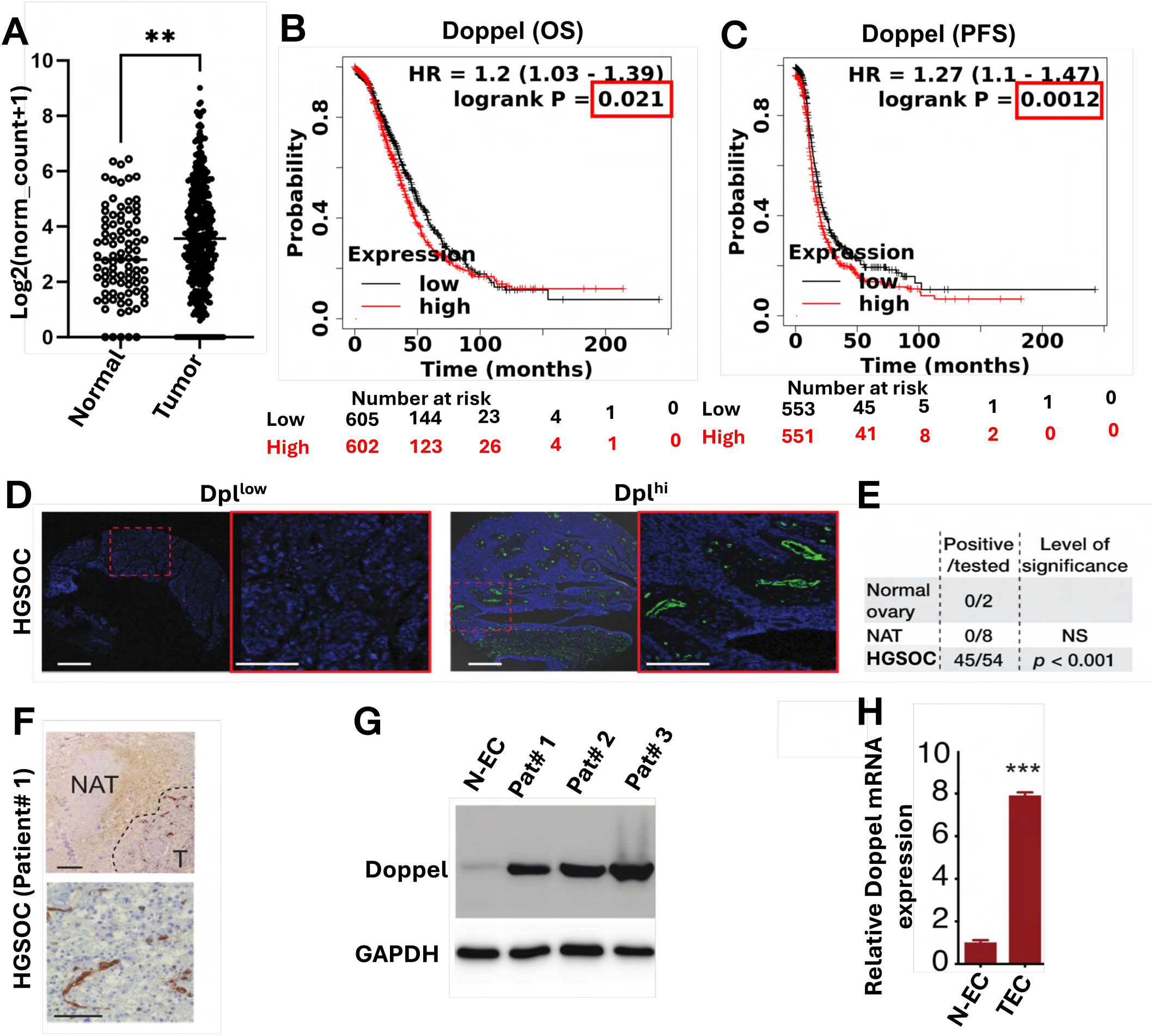
Doppel expression is elevated both in public and in-house ovarian cancer tumor samples. **A)** TCGA TARGET GTEx ovary RNA sequencing data comparison between serous ovarian cancer patients and healthy control tissue samples. Each dot indicates an individual subject. **B)** Kaplan-Meier overall survival analysis of public human serous ovarian cancer patients in the indicated groups with different Doppel levels. A total of 1207 serous ovarian cancer patients are included in this analysis. Cutoff value for low Doppel expression is 42. **C)** Kaplan-Meier progression free survival analysis of public human serous ovarian cancer patients in the indicated groups with different Doppel levels. A total of 1104 serous ovarian cancer patients are included in this analysis. Cutoff value for low Doppel expression is 40. **D and E)** Representative images of tumors from the HGSOC tissue microarray with low or high Doppel expression; 45 out of 54 tumor tissues were positive. **F)** IHC staining of Doppel in whole tissue section of HGSOC including tumor (T) and normal adjacent tissues (NAT). Doppel expression was exclusive to tumor vasculatures. **G and H)** Doppel protein and mRNA expression comparison between normal and tumor endothelial cells. Scale bars; 20 and 100 μm. *** p<0.001 and ** p<0.01. OS= Overall survival, PFS= Progression free survival, HR= Hazard ratio, Dpl= Doppel, HGSOC= High grade serous ovarian carcinoma, NAT= Normal adjacent tumor, T= Tumor, N-EC= Normal endothelial cells, TEC= Tumor endothelial cells.

### Doppel is specifically detected in the sera of ovarian cancer patients but not in benign or healthy female’s sera

Prion proteins are synthesized and processed through the secretory pathway, anchored to the cell membrane via a GPI anchor, and can be released into the extracellular space through proteolytic cleavage or via exosomes. For example, prion proteins are cleaved at tyrosine residue Y226 by the enzyme ADAM10 at the cell surface, releasing a soluble form into the extracellular space, which can then enter the bloodstream ^[15]^. After noticing the increased presence of Doppel protein in human ovarian carcinomas, we were intrigued to investigate whether Doppel is likewise discharged from the tumor tissues, as is seen in other secretory prion proteins. We detected Doppel concentration in a training cohort of archived sera samples of healthy and OC samples. In the training cohort, we detected serum Doppel in ovarian cancer patients (n=28) and healthy control (n=18). Doppel level was found to be significantly higher in ovarian cancer patients than in healthy control (14.24 ng/mL vs 1.06 ng/mL) and the receiver operating characteristic (ROC) curve between control and ovarian cancer patients confirmed the performance and applicability of Doppel as a sera marker **(Fig S1A and S1B)**. The statistics between control and patients in archived training cohort samples showed only two samples among 28 patients below cut-off value (4.77 ng/mL) **(Fig S1C),** further strengthening the applicability of Doppel as a sera biomarker.

To confirm the previous findings, we detected Doppel concentration in the 1^st^ validation set where we used an archived sera of 76 OC patients with stage and histopathology information, 10 triple negative breast cancer (TNBC) patients, and 18 healthy females using ELISA. Doppel level was significantly higher in sera of OC patients than their healthy counterparts (0.720 vs 6.653 ng/mL, p<0.0001); 72 out of 76 patients were positive for Doppel expression (cut off 2.67 ng/mL that include 99% of healthy samples) while only 1 out of 18 cases was positive for healthy female (**Fig. 2A**). Moreover, we didn’t see the differences between healthy and TNBC samples indicating Doppel specificity for OC diagnosis **(Fig 2A)**. The ROC curve analysis showed that Doppel can differentiate OC patients from normal individuals with AUC of 0.973, sensitivity of 0.91, specificity of 0.89 (**Fig. 2B**). Correlation of Doppel level with stages was also observed in HGSOCs patients (Stage I/II 5.28±0.96 ng/mL; n = 18 and Stage III/IV 9.64±1.15 ng/mL; n = 31 vs controls 0.72±0.19 ng/ml; *p*<0.0001 and *p*<0.01 between Stage I/II vs Stage III/IV) (**Fig. 2C**). The ROC curve analysis also showed that Doppel can differentiate early stage HGSOC patients from healthy individuals with AUC of 0.941, sensitivity of 0.94, specificity of 0.83 whereas the sensitivity and specificity for late stage HGSOC patients were 1.0 and 0.925, respectively (**Fig. 2D**). Interestingly, serum Doppel level was higher among EOCs of serous adenocarcinoma (7.020 ng/mL; n = 39), mucinous (10.780 ng/mL; n = 8), and endometroid (7.295 ng/mL; n = 7) than clear cell (3.114 ng/mL; n = 7) or other types (4.855 ng/mL; n = 15) (**Fig. 2E**). Together, the results from these two archived sera samples showed the potential of Doppel as a serological biomarker for the main subtypes of EOC detection and importantly, for early stage HGSOCs. However, the specificity of Doppel in benign conditions are unknown.

**Figure 2:**
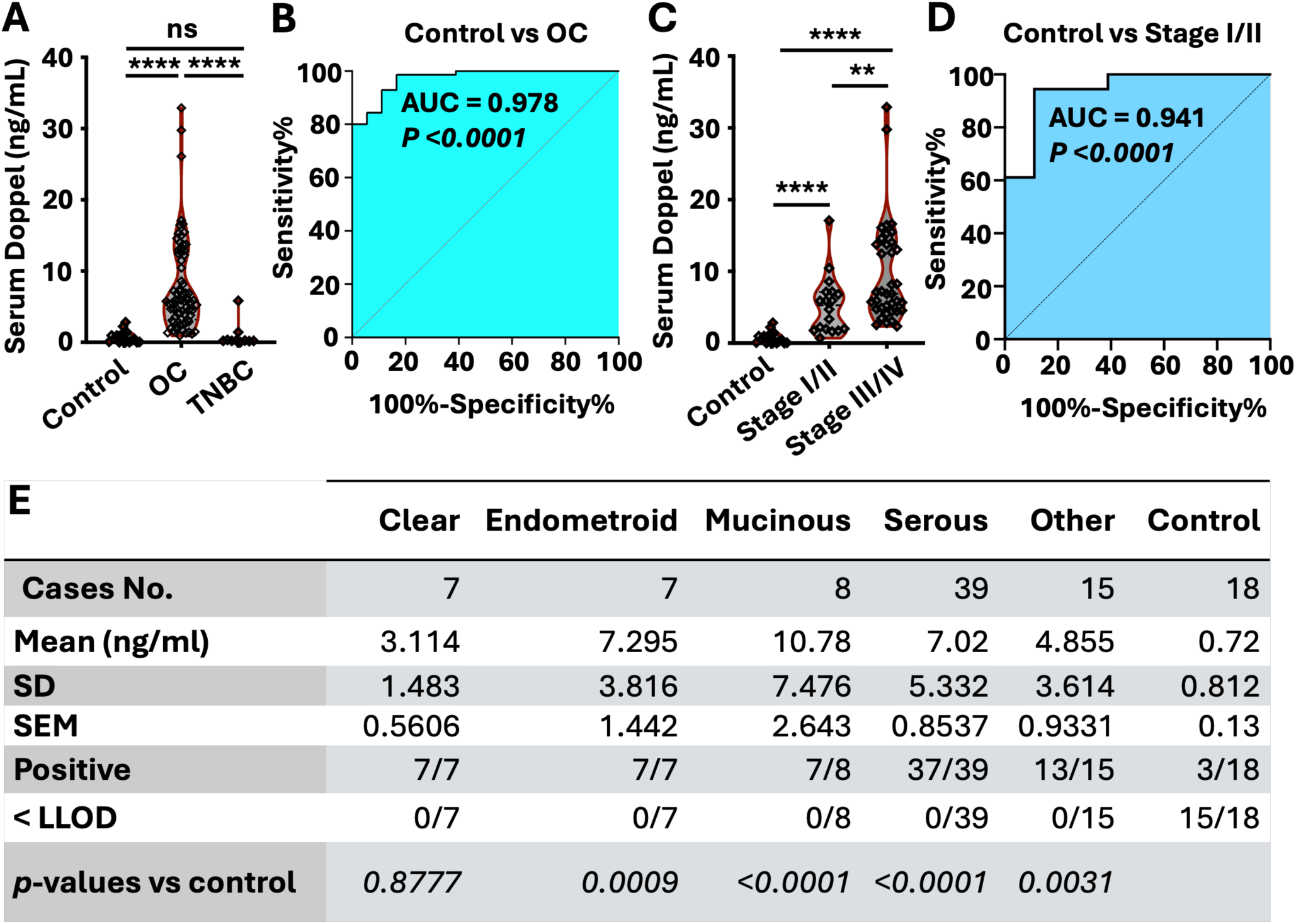
Serum Doppel level in patients with ovarian cancer subtypes. **A)** Measurement of serum Doppel level of healthy, OC and TNBC patients by ELISA. **B)** The ROC curve of Doppel between control and OC serum samples. **C)** Stagewise serum Doppel level in patients with HGSOCs compared with control subjects. **D)** The ROC curve of Doppel between control and HGSOC Stage I/II samples. **E**) Serum Doppel level based on ovarian cancer histology including epithelial ovarian cancer subtypes (clear cell carcinoma, endometroid, mucinous, and serous). ns= Not significant, ****= *p* <0.0001, OC = Ovarian cancer, TNBC= Triple-negative breast cancer, ROC= Receiver operating characteristic, AUC= Area under the ROC curve, HGSOC= High grade serous ovarian carcinoma.

To confirm the specificity of Doppel as EOC biomarker, we used a 2^nd^ validation cohort which included healthy, EOC and benign patients. We recruited age, race, and ethnicity-matched 17 control females and treatment-naïve 22 cases (**Table S1**), out of the 22 cases, 7 are diagnosed as benign, 15 as EOCs. We detected the Doppel level in control, benign and EOC subjects and the results showed EOC subjects Doppel level significantly higher than control and benign subjects (**Fig 3A**), whereas no statistical differences are seen between control and benign subjects clearly indicating the Doppel specificity in EOCs. CA-125, a widely used FDA approved marker for OC detection, sometimes fails to detect OCs due to its lower limit of detection (LLOD; <35 IU/mL) and the occurrence of false positive results in benign tumor due to specificity issue ^[16]^. We compared our Doppel sera findings of EOC and benign subjects with CA-125, a pairwise comparison revealed that CA-125 level was lower than its cutoff value (35 IU/mL) ^[17]^ in two EOC samples (33 and 12.7 IU/mL) **(Fig 3B)** and higher in two benign samples (158 and 312 IU/mL) **(Fig 3C)**. However, Doppel level was consistently higher or lower than its cutoff value (2.67 ng/mL) in EOC **(Fig 3B and 3C)**. These findings collectively validated our exploratory sera Doppel results and provide proof of concept of Doppel usefulness as OC biomarker over CA-125 to distinguish and identify benign and high-grade ovarian cancer patients.

**Figure 3:**
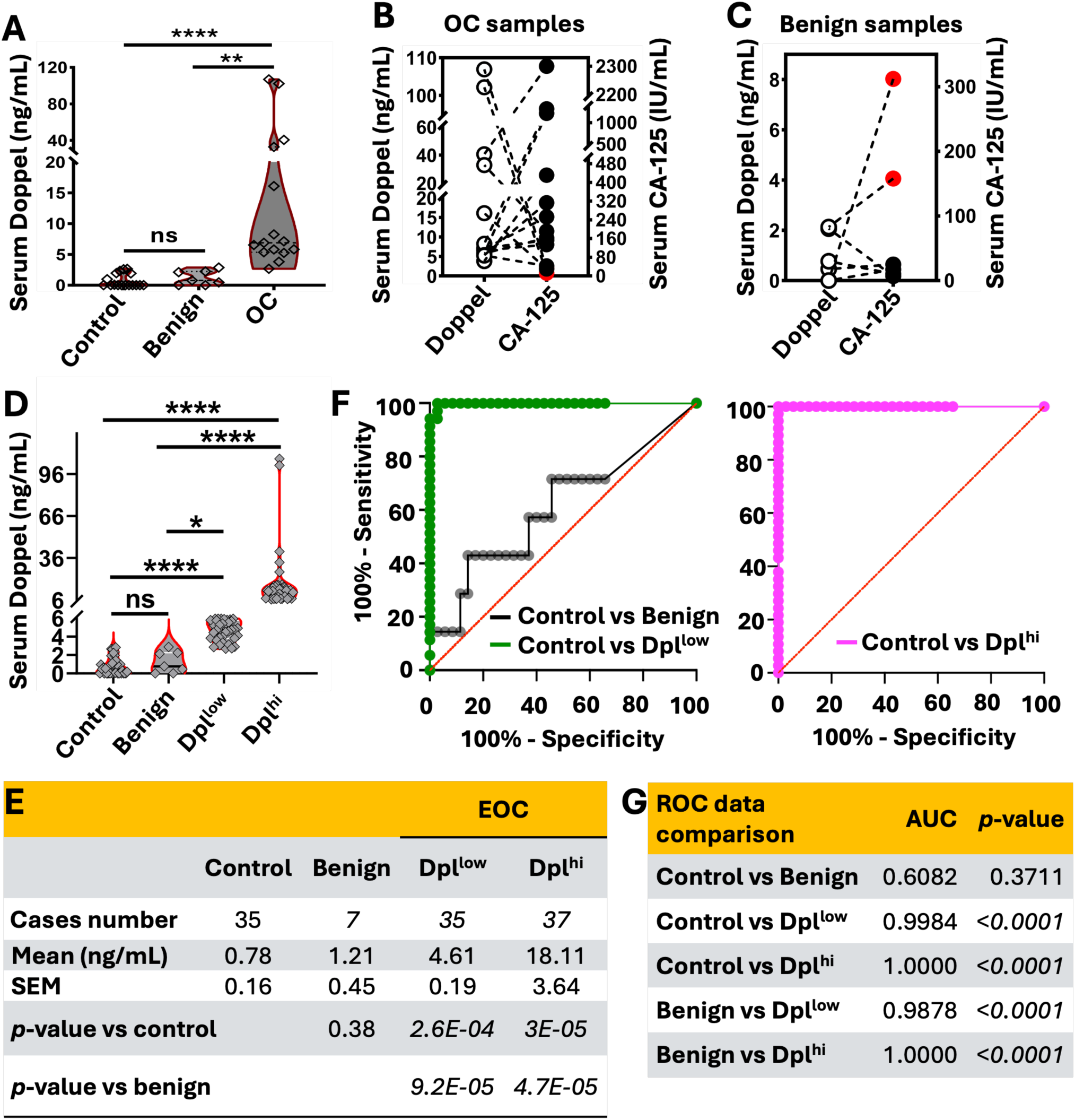
Doppel level distinguishes cancerous from non-cancerous patients. **A)** Comparison of serum Doppel level of OC patients with benign and healthy control samples. **B)** Pairwise correlation of serum Doppel with serum CA125 level in OC ovarian cancer patients. **C)** Pairwise correlation of serum Doppel with serum CA125 level in patients with benign tumors. **D, E)** Level of Doppel in healthy, benign and EOC patients that are stratified into Dpl^hi^ and Dpl^low^ groups. **F)** The ROC curve between control vs benign, control vs Dpl^low^ and control vs Dpl^hi^ groups. **G)** The AUC and *p*-value of ROC plots between control vs benign, control vs Dpl^low^, control vs Dpl^hi^, benign vs Dpl^low^, benign vs Dpl^hi^ groups. Serum and ascitic Doppel level are measured by ELISA. ns= Not significant, *= *p*<0.05, **= *p*<0.01, ****= *p*<0.0001. EOC= Epithelial ovarian cancer, ROC= Receiver operating characteristic, AUC= Area under the ROC curve.

### Stratifying ovarian cancer based on Doppel level in patient’s sera

Profiling of patients with high and low Doppel-level may be mapped to identify the significance of Doppel-axis in ovarian cancer progression. Based on the serum Doppel level, combining the EOC samples from our 1^st^ and 2^nd^ validation sets (n = 72), we proposed a Doppel-index and classified them into Doppel-high (Dpl^hi^) or Doppel-low (Dpl^low^) groups. If the serum Doppel level is >6 ng/mL, we categorized them into Dpl^hi^. If the serum Doppel level is higher than the cutoff value (>2.67 ng/mL) but <6 ng/mL, we categorized them into Dpl^low^. Serum Doppel exhibited a significantly elevated level in both Dpl^hi^ (18.11 ng/mL; n=37) and Dpl^low^ (4.61 ng/mL; n=35) in comparison to control and benign groups **(Fig 3D and 3E)** and all found higher than LLOD. Moreover, in benign cases Doppel level was similar to control (1.2 ng/mL in benign and 0.74 in control) with no significant differences **(Fig 3D and 3E)**. The ROC curve achieved high accuracy for EOCs detection for both Dpl^low^ groups with an AUC of 0.99 and Dpl^hi^ groups with an AUC of 1.00 in comparison to control (**Fig 3F and 3G**). Similar results were obtained for both Dpl^low^ and Dpl^hi^ groups when the data were compared with benign groups (**Fig 3G**).

### RNA sequencing of patient-derived organoids from Doppel-high versus low groups revealed induction of epithelial-mesenchymal transition in ovarian cancer cells

The number of integrated and exploratory biomarkers in ovarian cancer detection is increasing, reflecting a potential clinical significance. Nevertheless, despite the anticipation surrounding their addition, many of these biomarkers frequently lack evidence of their involvement in the progression of ovarian cancer. The poor prognosis of ovarian cancer is associated with malignant ascites formation; fluid developed in OC patient’s peritoneal cavity where cancer cells survive. Malignant ascites is common in ∼75% of patients upon initial diagnosis and prevalent among patients who exhibit resistance to chemotherapy ^[18]^. This indicates that analyzing the molecular composition of malignant ascites in OC could offer valuable insights for clinical monitoring. To this end, we checked Doppel level in eight patient’s sera and ascites that were collected retrospectively (**Table S2**). All the patients were positive for Doppel detection based on our cutoff value (2.67 ng/mL) and the data revealed a strong correlation between sera and ascites Doppel level **(Fig 4A)**, suggesting a role of Doppel in ascitic transformation. However, 4 out of 8 patients had high Doppel expression in their sera and ascites that allowed us to categorize them into Dpl^hi^ and Dpl^low^ groups **(4A)**.

**Figure 4:**
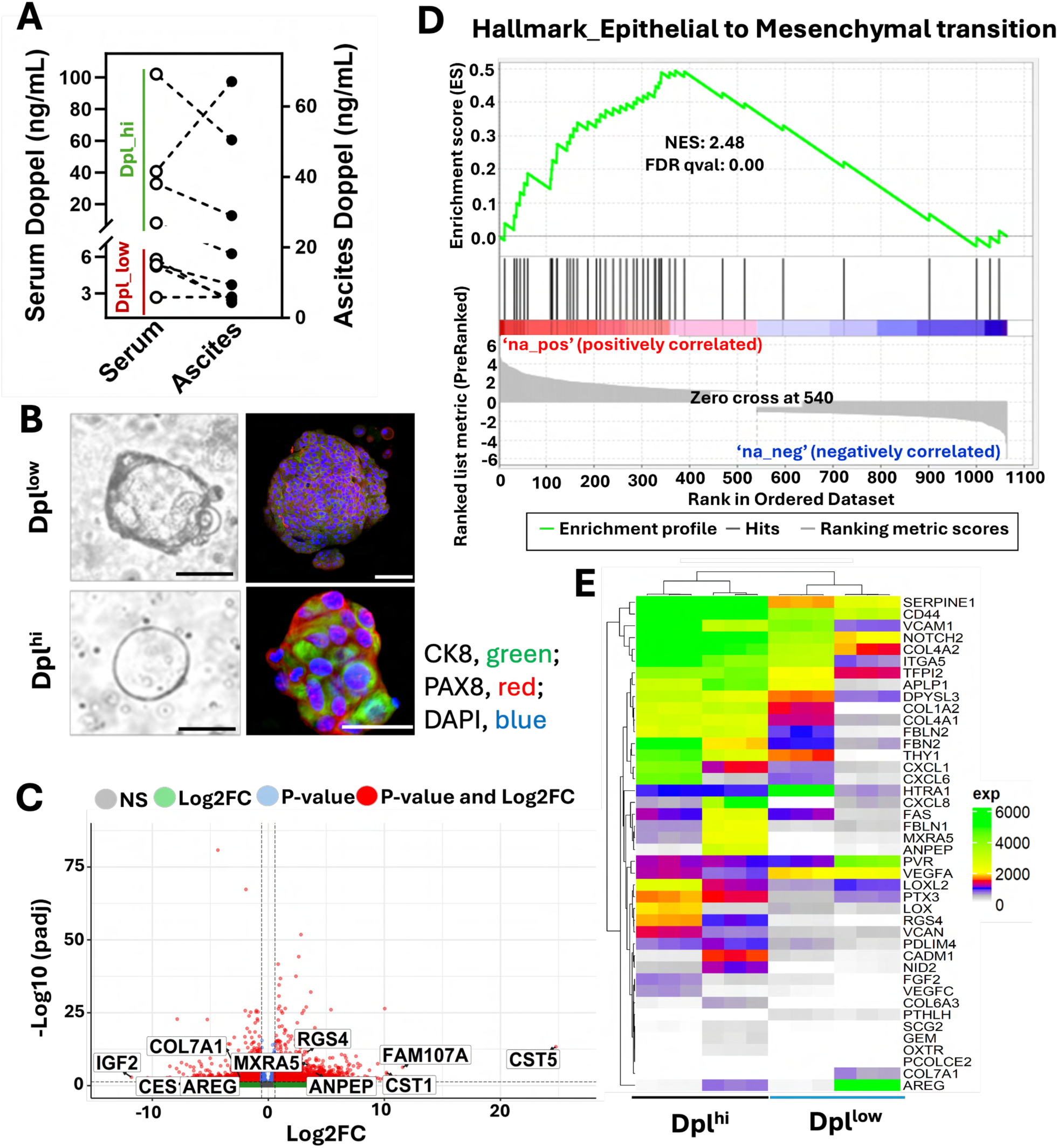
Doppel expression induced EMT in ascites-derived organoids. **A)** Pairwise comparison of serum Doppel with ascitic Doppel level in ovarian cancer patients. **B)** Representative bright field (left, Scale bar: 200 μm) and immunofluorescence (right, Scale bars: 200 and 50 μm) images of organoids established from patient’s ascites with Dpl^low^ and Dpl^hi^ level. OC specific markers (CK8, PAX8) were used to confirm the establishment of organoids. **C)** Volcano plot of genes in Dpl^hi^ organoids compared to Dpl^low^ organoids; y axis represents the negative Log 10 of p adjusted value; x axis represents the Log2 fold changes between groups. **D)** GSEA analysis showing the enrichment of EMT hallmark pathway in Dpl^hi^ organoids. **E)** Heat map showing the EMT associated genes in Dpl^hi^ versus Dpl^low^ organoids. Normalized genes expression values are used. NES= Normalized Enrichment Score, GSEA= Gene set enrichment analysis, EMT= Epithelial mesenchymal transition.

To understand the function of Doppel in malignant ascites, we chose four ascites samples from two each from Dpl^hi^ and Dpl^low^ group with similar histology (serous and mucinous) type (**Table S2**). To uncover the phenotypical differences between Dpl^hi^ and Dpl^low^ groups, we developed organoids from ascites derived cells following standard protocol ^[19]^ ^[20]^. Representative images of organoids show the morphological features of established organoids and expressions for OC markers: CK7 and PAX8 **(Fig 4B)**. We performed RNA sequencing using mRNA collected from ascitic organoids and identified a total of 1138 (549 up and 588 down) differential expressed genes (DEGs) between Dpl^hi^ and Dpl^low^ groups based on fold change (−1≥Log2FC≥1), padj≤0.05 and count ≥10. Volcano plot showing all the genes including up and down in Dpl^hi^ and Dpl^low^ groups **(Fig 4C)**. To identify the significant biological processes of significant DEGs between Dpl^hi^ and Dpl^low^ groups we performed hallmark GSEA and identified epithelial-mesenchymal transition (EMT) associated with Doppel high group **(Fig 4D)**. The heatmap depicting 42 genes related to epithelial-mesenchymal transition (EMT) highlights the consistent disparities between Dpl^hi^ and Dpl^low^ groups **(Fig 4E)**.

### Single-cell RNA sequencing of whole ascites identifies a strong correlation between protumorigenic macrophages and patients with Doppel-high levels

In ascites, disseminated OCs have to survive in a peritoneal environment constituted by lymphocytes, macrophages, natural killer (NK) cells, fibroblasts, mesothelial cells, as well as cytokines and chemokines secreted by these cells for a successful transcoelomic metastasis^[21]^. Although the role of the immune microenvironment in peritoneal cavity is indisputable to the spread of OC, the molecular network regulating the crosstalk between the disseminated tumors and immune cells in peritoneal cavity is highly unknown. Therefore, we tried to explore the role of Doppel as a novel regulator of ascitic immune microenvironment that would be imperative to understand its function with OC progression. We performed single-cell RNA sequencing (scRNAseq) of organoid-matched whole ascites-derived cells. After filtering the low-quality cells, integration of individual patients was performed for future integrated analysis. Figures **S2A** and **S2B** confirm the successful integration and cell-cycle analysis result and indicate no cell-cycle heterogeneity are present in our dataset after integration. Using the uniform manifold approximation and projection (UMAP) method, we discerned 9 primary cell clusters, encompassing both immune and non-immune cell types **(Fig 5A)**, based on marker genes shown in figure **S2C**. Immune cell types are T cells, B cells, NK cells, Dendritic cells, macrophages and plasma cells, whereas non-immune cells are mainly epithelial-originated tumor cells and small portions of stromal cells. A portion of cells only express PTPRC **(Fig S2C)**, present in both groups and cluster with T cells subcluster **(Fig 5A)**, indicating the complex ecosystem of immune activity in ascites.

**Figure 5:**
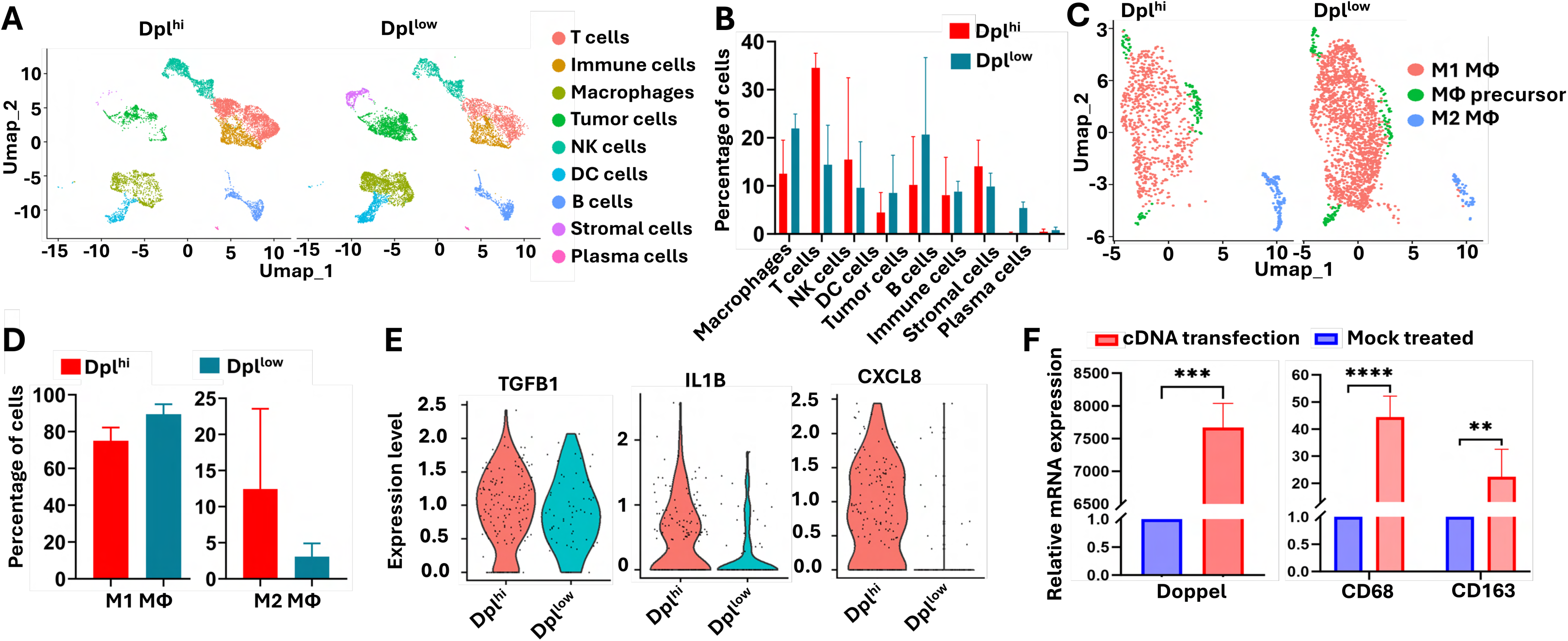
Doppel expression is associated with M2 macrophage phenotypes in malignant ascites. **A)** UMAP plot of scRNA-seq showing the distribution of different cell populations in Dpl^hi^ and Dpl^low^ ascites. **B)** The percentage of cells, in primary clusters, present in each ascites. **C)** UMAP plot showing the distribution of macrophage cell populations in Dpl^hi^ and Dpl^low^ ascites. Secondary analysis was performed on macrophage cluster. **D)** Bar chart showing the percentage of antitumorigenic (M1) and protumorigenic (M2) macrophages, in secondary clusters, present in each ascites. **E)** Violin plot showing the key M2 macrophage associated marker genes in secondary macrophage M2 cluster in Dpl^hi^ and Dpl^low^ ascites. **F)** Left, Doppel mRNA expression in mock and Doppel cDNA transfected monocytes (U937). Right, CD68 and CD163 mRNA expression in mock and Doppel cDNA transfected monocytes (U937). Error bar represents mean ± SD. **= *p*<0.01, ***= *p*<0.001, ****= *p*<0.0001. UMAP= Uniform Manifold Approximation and Projection, scRNA-seq= Single cell RNA sequencing.

To match our scRNAseq findings with the organoid model, we identified the DEGs between Dpl^hi^ and Dpl^low^ groups of epithelial-originated tumor cells and compared them with DEGs (top 50 up- and down-regulated) of same group as that of the organoid model. The results indicated that genes found in epithelial-originating tumor cells exhibited a similar pattern to that observed in the organoid model across the various groups, except for ZBTB16 **(Fig S2D)**, further reinforcing the significance of the organoid findings. Percentage of cells present in each group **(Fig 5B)** indicated the dynamic changes of immune populations and the consistent changes of macrophages and T cells population prompted us to subcluster them to explore their distinct function. Our results didn’t show any significant changes among identified T cell subpopulations between Dpl^hi^ and Dpl^low^ group **(Fig S2E, S2F and S2G)**.

Next, we performed unsupervised clustering of macrophages and identified 3 clusters including M1 and M2 macrophages **(Fig 5C)**. We noted the coexistence of both M1 and M2 functional phenotypes, underscoring their intricate roles, which align with findings from prior research ^[22]^. The expression of marker genes within macrophage subclusters also demonstrates the simultaneous presence of both M1 and M2 phenotypes **(Fig S2H)**. However, based on higher CD163 expression **(Fig S2H)**, we successfully identified M2 macrophages activity which is higher in Dpl^hi^ and Dpl^low^ group **(Fig 5D)**, conclusively underscoring their significant contribution within the ascitic tumor microenvironment (TME) of Dpl^hi^ group. The M2 macrophages cluster preferentially expressed immunosuppressive genes in Dpl^hi^ group including TGFB1, IL1B and CXCL8 **(Fig 5E)**. To validate our scRNAseq findings of M2 macrophage characteristics, we overexpressed Doppel on monocytic U937 cells and confirmed that Doppel significantly induces CD68/CD163 expression, indicating M2 phenotypes **(Fig 5F).** Overall, using two distinct and advanced *in-vitro* models, we have pinpointed the involvement of Doppel in fostering cancer cell aggressiveness and immune suppression.

### Elevated Doppel level in the sera and ascites correlates with the dissemination of ovarian tumor cells into the circulation

If Doppel regulates EMT of OC cells and ascitic TME of OC patients by inducing pro-tumorigenic macrophages, we hypothesize that Doppel controls the dissemination of ovarian cancer into the circulation. To detect circulating tumor cells (CTCs) in blood of Dpl^hi^ and Dpl^low^ patients, we devised an in-house microfluidic chip model that was coated with Dpl-Ab **(Fig 6A)** and identified CTCs based on their expression of DAPI, EPCAM and cytokeratin, while noting the absence of CD45 expression in CTCs **(Fig 6B)**. We detected and counted the Doppel expressing CTCs from fresh whole blood collected from 7 Dpl^hi^ and Dpl^low^ patients. As in the validation experiments, the anti-EPCAM antibody was used as the capturing antibody and the fluorophore-labeled-cytokeratin as the target label. Overall, Dpl-Ab coated chips showed similar efficiency of TC capturing as with EPCAM-ab coated chips (**Fig. 6C**), which suggests CTCs exclusively express Doppel. Thus, we termed the Dpl-Ab captured CTCs as ^Dpl+^CTC. ^Dpl+^CTC were significantly higher in the blood of Dpl^hi^ than Dpl^low^ patient groups **(Fig 6D)**, strongly indicating the association of Doppel with OC dissemination into the circulation. Next, we performed pairwise comparison of serum Doppel level with CTC counts in the blood. Despite some variability, the majority of Dpl^hi^ patients are associated with higher counts of CTCs in blood **(Fig 6E)**. We further performed the pairwise comparison between ascites Doppel level and CTC counts in blood. Although having some variability, a correlation was established between ascites Doppel and the number of ^Dpl+^CTC in the blood **(Fig 6F)**. These findings collectively established a novel link between serum/ascites Doppel levels and ovarian cancer dissemination into the circulation.

**Figure 6:**
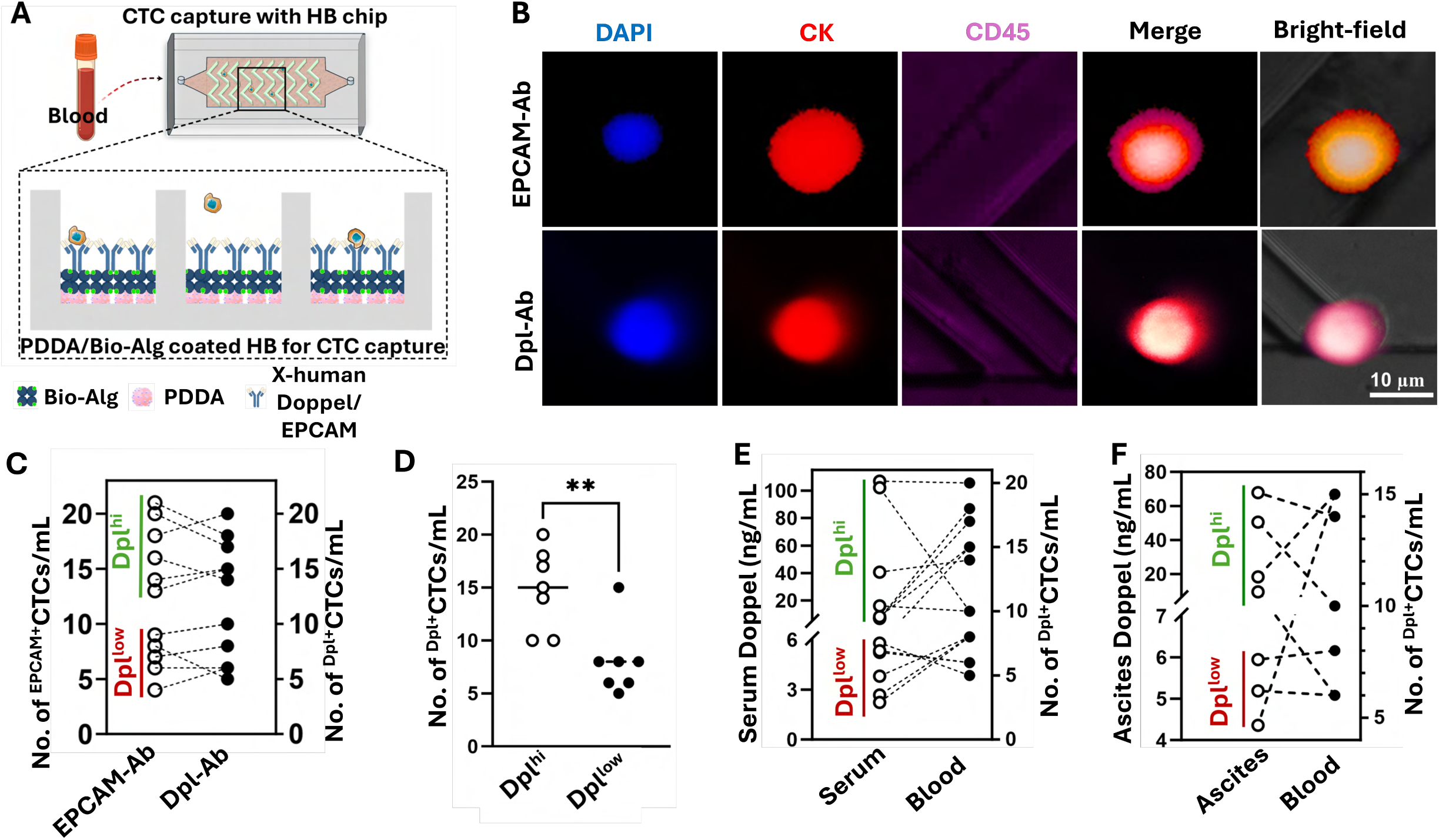
Serum and ascitic Doppel level correlate with circulating tumor cells in blood of EOC patients. **A)** Schematic of microfluidic chip used to isolate EPCAM and Doppel expressed CTCs. **B)** Representative immunofluorescence images of markers expression in isolated CTCs. DAPI^+^, CK^+^, CD45^-^ cells are considered CTCs. Upper panel: CTCs captured by EPCAM antibody- coated chips and lower panel: CTCs captured by Doppel antibody-coated chips. **C)** Pairwise comparison of isolated ^EPCAM+^CTCs and ^Dpl+^CTCs in the patient’s blood. **D)** No. of ^Dpl+^CTCs isolated from 14 EOC patient’s whole blood that are stratified into Dpl^hi^ and Dpl^low^ groups. **E)** Pairwise comparison of serum Doppel level with whole blood ^Dpl+^CTCs. **F)** Pairwise comparison of ascites Doppel level with whole blood ^Dpl+^CTCs. DAPI= 4’,6-diamidino-2-phenylindole, CK= Cytokeratin, Dpl= Doppel, CTC= Circulating tumor cells, Ab= Antibody, EPCAM= Epithelial cell adhesion molecule. **= *p*<0.005.

## Discussion

Taking into account the limitations of existing diagnostic modalities, in this study, we uncover for the first time that Doppel, a testis-specific prion-like protein, aberrantly expresses in OC tissues– mainly within their vasculatures and it’s soluble form is detectable in patient’s sera and ascites. Doppel can serve as a biomarker for OC, effectively distinguishing EOC patients, mostly early stages of high-grade serous cases from those with benign conditions and controls. Clustering the EOC patients with Dpl^hi^ and Dpl^low^ groups, we also identify how Doppel may influence ascitic TME and cancer cells to disseminate into the circulation. We demonstrate that Doppel influences EMT pathway by leveraging the function of M2 macrophage in ascitic TME, potentially resulting in a substantial increase in CTCs among OC patients.

Our study has two-fold implications. First, the specificity and sensitivity of Doppel (AUC of 0.973 for Dpl vs control) surpasses that of CA125; CA125 was lower than its cutoff value (>35 IU/mL) in at least 20% of ovarian cancer cases in our cohorts as well as reported analyses ^[23]^. Further comparison between Doppel and CA125 for identifying benign conditions clearly indicates the superiority of Doppel over CA125. Importantly, we found elevated Doppel level in a number of matched ascites of OC patient and a linear relationship between sera and ascitic Doppel level, indicating the involvement of Doppel in ascitic TME. Our data represent a clinical significance, as ascites is found in over 90% of patients diagnosed with stage III/IV OC, and the correlation between ascites and disease progression is notable ^[24]^. In addition, ascites is a significant adverse predictor of adverse outcome in recurrent disease ^[23]^. The observation that Doppel is not expressed in healthy ovaries, or any other reproductive organs of females suggest that this marker is a bona fide marker of neoplastic transformation. We identified serum Doppel as early as Stage I/II of EOCs and significantly stepwise increase among HGSOCs, suggesting that Doppel expression would be important to detect the early stages of HGSOCs. Acquiring patient samples from multiple institutions also enable us to account for ethnic and geographical diversities. However, OC encompasses various invasive ovarian carcinomas originating from diverse tissues, including high-grade serous, clear cell, endometrioid, and mucinous subtypes. HGSOC arises from the distal fallopian tube epithelium, but Doppel is not a specific marker for fallopian tube biology. Thus, how early the transformed epithelial cells express Doppel would be interesting to study. Understanding of Doppel-biology may deepen our insight into HGSOC’s origin and early evolution.

Second, using the muti-omics approach with patient-matched ascites and blood samples, we found that soluble Doppel level creates an immunosuppressive ascitic TME and correlates with EOC aggressiveness. In the primary tumor tissues, as we have seen Doppel expression in tumor vasculatures (**Fig 1**), this protein is known to regulate neo-angiogenesis where Doppel-VEGFR2- axis plays the most crucial role ^[14]^. However, in malignant ascites, we found a causative role of Doppel with EMT-phenotype. The top two EMT related abundant genes in Dpl^hi^ group, serpin family e member 1 (SERPINE1) and CD44, were reported to induce EMT in ovarian cancer ^[25]^ ^[26]^. Ascites has both cellular and acellular components, where acellular components can induce EMT ^[27]^.These findings suggest that soluble Doppel in ascites may prime cancer cells, thereby significantly contributing to the promotion of EMT. Further investigation into the interaction between EMT genes and soluble Doppel could provide deeper insights into the mechanisms driving OC EMT in this specialized compartment.

Single cell RNAseq of ascites samples identified a pro-tumorigenic M2 macrophage phenotype in Dpl^hi^ expressing groups that coincide with *in-vitro* data. Consistent with earlier findings ^[28]^ ^[29]^ ^[30]^, we observed the elevated activity of transforming growth factor (TGFB1), interleukin-1B (IL1B) and chemokine (C-X-C motif) ligand 8 (CXCL8) correlating with increased Doppel activity in M2 macrophage cell population. Analysis of T cells in clinical ascites revealed no significant differences between the Dpl^hi^ and Dpl^low^ groups. Our results are not surprising, considering the fact that protumor effects in peritoneal fluid are not influenced by T lymphocytes ^[31]^. Finally, we showed a link between CTC dissemination and Doppel expression. We found that the number of CTCs increased in Dpl^hi^ OC group. This finding is interesting because we captured CTCs based on their ability to express Doppel. Doppel is known as a surface marker, and we predominantly observed Doppel expression in the tumor endothelial cells of primary tumors. However, we have never tested whether Doppel expresses in transformed cancer cells. In support of this study, we found that Doppel was exclusively detected patient-derived EOC organoids. Thus, the detection of ^Dpl+^CTC is not surprising in these transformed EOC cells. Also, previous studies demonstrate that CTCs with EMT-phenotype are common in OC patients ^[32]^. Since Doppel induces EMT and aids CTC dissemination in our patient-matched cohorts, we posit that Doppel-indued EMT may prime OC cells to disseminate into the circulation.

Although we provided proof-of-concept for Doppel as an OC biomarker, we have not directly assessed the association of Doppel expression with patient survival and mortality in our sample cohort. Future work should determine the effect of higher Doppel expression with patient outcome using large cohort of patients across the entire spectrum of the disease (e.g., benign, early, late stages as well as rare ovarian cancer subtypes). In addition, it will be interesting to check whether Doppel can be used to identify recurrence in patients with OC by using prospective samples, as the sensitivity of CA-125 for recurrent EOCs varied widely (56-94%)^[33]^. To identify the association of Doppel in the ascitic TME, we used patient-matched samples combined with advanced *in vitro* model and sequencing techniques. Although we performed analyses in small sample sizes, these combined approaches significantly advance our understanding of Doppel- biology in inducing EMT or manipulating the ascitic immune microenvironment of OC. However, the activity of Doppel in the ascitic TME will need to be assessed systematically using appropriate models. While the association of serum and ascitic Doppel level with blood-derived ^Dpl+^CTCs relates to the role of Doppel in OC dissemination into the circulation, the significance of ^Dpl+^CTCs may be enormous. For instance, analysis of ^Dpl+^CTCs could have potential for metastatic propensities. Longitudinal monitoring of serum Doppel or ^Dpl+^CTCs in the surveillance settings could improve a patient’s chances of survival.

Collectively, our findings identified Doppel as a novel biomarker for OC diagnosis and highlight a mechanism of Doppel-axis in OC ascitic TME. Our other major contribution is the detection of CTC based on Doppel expression, ^Dpl+^CTC. We showed two platforms, soluble Doppel in sera/ascites and ^Dpl+^CTC in whole blood, to detect subsets of EOCs that may be further developed for clinical validation. Also, ascites offers valuable insights into the primary tumor’s progression to metastasis. Thus, targeting and studying Doppel-axis in ascites of EOCs may advance our understanding of key signaling pathways and identify new therapeutic strategy.

## Materials and methods

### Collection of clinical samples

We collected clinical samples from three different cohorts. For the training cohort, sera collected from 28 ovarian cancer patients and 18 healthy female subjects were analyzed from an archived cohort ^[34]^. For the 1^st^ validation cohort, sera collected from 76 ovarian cancer patients, 10 triple negative breast cancer (TNBC) patients, and 18 healthy females were analyzed. Serum samples were collected at the Korea University Anam Hospital from 2006 to 2020 from patients who underwent conservative, comprehensive or staging surgery for ovarian cancer. Preoperative imaging procedures (ultrasonography and computed tomography) were used to identify the presence of ovarian cancer with stages from I to IV. In the 2^nd^ validation cohort, 22 ovarian cancer and 7 benign cancer patients were recruited, and specimens were collected during surgery at Texas Tech University Health Sciences center El Paso under Institutional Review Board-approved protocols (IRB#: E21144) between October 2021 and December 2023. The disease stages of recruited patients were clearly defined in IRB protocol, based on 2018 International Federation of Gynecology and Obstetrics classification system. **Patient inclusion criteria:** patients between 35- 85 years of age with no severe autoimmune disease or previous history of cancer were included. Ovarian cancer cohorts comprise benign, borderline (BL) cases, type I cases and type II cases, and stage I-IV. Type I cases and type II cases were clearly defined in IRB protocol. **Patient exclusion criteria:** Patients with previous history of familial ovarian cancer and ages below 35 years of age were excluded from the analysis. Patient-matched control sera were collected from 17 healthy females. **Control inclusion criteria:** 35-85 years of age control subjects with no prior history of seizure disorder, head trauma (within less than one year), cancer, CNS tumor, hepatic cirrhosis, connective tissue disease (ie: lupus, rheumatoid arthritis, etc), congenital heart disease was considered as healthy control. **Control exclusion criteria:** following exclusion criteria were maintained while recruiting control subjects- women under the age of 35, planned surgical intervention, decline phlebotomy, self-reported previous bilateral oophorectomy, subjects diagnosed with cancer. Blood and ascites were drawn and drained respectively by interventional radiologist and transferred for further processing in appropriate containers. The demographics of these patients are summarized in **Table S1**.

### Enzyme-linked immunosorbent assay (ELISA)

Serum and ascites Doppel level was measured using Human Prion-like protein Doppel ELISA Kit, 96-Strip-Wells (MBS76513, MyBioSource, CA) according to the manufacturer’s instructions. Briefly, we aliquoted 100 µL of standards and 100 µL of diluted samples into pre-coated plate, incubated the sealed plate at 37°C for 90 minutes, and then washed twice with wash buffer. 100 µL biotin-labeled antibody solution, 100 µL HRP-streptavidin conjugate and 90 µL TMB substrate were added sequentially to each well. Between each step, incubation and washing steps were performed following protocols. At the end of the experiment, we stopped the reaction with 50 µL stop solution, and immediately measured the absorbance at 450 nm. The following formula was used to measure the sample concentration: (the relative O.D. 450) = (the O.D. 450 of each well) - (the O.D. 450 of blank well). A standard curve was made by plotting the relative O.D. 450 of each standard solution vs the respective concentration of the standard solution. To fit the concentration of Doppel within standard, we made 2 dilutions of each sample (1:20 and 1:40) with triplicates. The final concentration was measured by multiplying the dilution factor to the concentration from interpolation and calculated the average result considering both dilution and triplicate samples of each. For each sample, we conducted four replicates and calculated the concentration as the average of these measurements.

### Bioinformatics analysis

Human ovarian cancer Doppel gene-expression TCGA data (n=419) and normal Doppel gene-expression ovary data (n=88) were downloaded from Xena Browser (https://xenabrowser.net/) and Doppel expression differences between these two groups were calculated in GraphPad Prism 10. Overall and progression free survival associated with Doppel expression in ovarian cancer were performed in Kaplan-Meier Plotter (https://kmplot.com/analysis/), a web-based survival analysis platform of various cancers. For survival analysis, we choose PRND gene (affymetric ID 222106_at) with median survival option and choose serous histology to determine the overall and progression free survival associated with Doppel expression.

### Immunofluorescence, immunohistochemistry, western-blot and quantitative PCR

Tissue microarray was purchased from US Biomax, Inc., which included 54 cases of HGSOC, 8 normal adjacent tissues (NAT), and 2 ovary tissues. Previously published protocols from our lab were used to perform immunofluorescence, immunohistochemistry, tumor endothelial cells isolation, western-blot and qRT-PCR ^[14]^. Each experiment is conducted as biological triplicate for western-blot and qRT-PCR. All the antibodies, reagents and primers used were mentioned in our previously published article except primers for Doppel, CD68 and CD163 ^[14]^. Following primers are used to quantify Doppel, CD68 and CD163 gene-expression by qRT-PCR (hDoppelF- GATGGCATCCACTACAACGGCT, hDoppelR- GTTGTCTGGCTTCTGGAACTCC, hCD68F- CGAGCATCATTCTTTCAGCT, hCD68R- ATGAGAGGCAGCAAGATGGACC, hCD163F- CCAGAAGGAACTTGTAGCCACAG, hCD163R- CAGGCACCAAGCGTTTTGAGCT).

### Preparation of serum, ascites and development of organoid model

Freshly collected blood samples in K2EDTA vacutainer tube were centrifuged for 20 minutes at 4^0^C at 1000×g and clear supernatant (serum) were collected and stored aliquoted at −80^0^C for future experiment. Malignant ascites samples were collected fresh from ovarian cancer patients undergoing surgery. The ascitic fluid was centrifuged at 1600× g for 10 minutes at 4°C and the supernatant was collected and stored at −80^0^C for future experiments. To remove the red blood cells (RBC), pelleted cells were resuspended and incubated with diluted RBC lysis buffer (sc- 296258, Santa Cruz, TX) for 10 minutes at 4^0^C and centrifuged down at the RBC cleared cells. If RBC was still visible in the pellet, a second round of RBC lysis was performed. The resulting pellet washed with cold PBS and manual cell counting was performed. The samples were divided into many portions based on cell counting. For organoid culture, freshly pelleted cells resuspended and cultured following modified protocols adapted from published article ^[19]^ ^[20]^. A separate article describing the detail protocol of organoid development from ascites will be published.

### RNA isolation and bulk RNA sequencing analysis

RNA was extracted from mature organoid using combined TRIzol (15596026, Thermo Fisher Scientific, MA) and RNeasy Mini Kit (74104, Qiagen, MD) following the manufacturer’s guidelines. The quality of RNA was determined by Nanodrop absorbance (260/280 ratio ≥1.8) and RNA integrity value (≥7) analysis. Bulk RNA-seq was carried out by Novogene using the Illumina NovaSeq6000 platform. For each organoid we conducted three biological replicates. Adaptor trimming and quality of raw sequencing files were evaluated using FastQC software (0.12.1), considering metrics such as sequence quality scores, sequence duplication, and adapter content to determine if filtering was needed prior to genome mapping. Clean reads were then aligned to the human GRCh38 reference genome using the HISAT2 (2.2.1) graph-based read aligner. The aligned reads were assembled into transcripts or genes using StringTie software (2.2.0) considering the Fragments Per Kilobase of transcript sequence per Millions base pairs sequenced (FPKM) approach. Differential gene expression analysis was performed on raw counts with the R statistical package DESeq2 (3.19). Genes exhibiting a Log2 fold change (Log2FC) of ≥1, p_adj ≤ 0.05 and count≥10 were deemed significantly upregulated, while those with a Log2FC ≤ 1, p_adj ≤ 0.05 and count≥10 were considered significantly downregulated. Gene set enrichment analysis, using hallmark gene set, of significantly dysregulated genes was performed by GSEA (4.3.3) software. The heatmap of targeted pathway was generated in ‘R’ environment (4.3.2) using ComplexHeatmap (2.18.0) package.

### Single-cell RNA sequencing (scRNAseq) analysis

Preprocessed ascitic cells in cell storage media were sent to Novogene for scRNAseq library preparation, sequencing, and unique molecular identifier (UMI) count. The quality of each sample was measured by viability assay and samples having viability over 80% were used for scRNAseq library preparation. scRNAseq libraries of 5000 cells/ sample were generated by 10X genomics chromium next gem single cell 3ʹ v.3.1 and sequencing was performed by Illumina NovaSeq6000 platform with 50,000 reads/cell targeting 90G of total data for each sample. Reads from scRNAseq were initially processed for each sample using Cell Ranger (10X Genomics, version 7.1.0) with default settings. In summary, this involved collapsing UMIs, aligning reads to the human reference genome (GRCh38), counting UMIs, and performing initial quality control. Filtered feature barcode matrix, Cell Ranger output, files were imported to ‘R’ environment (4.3.2) and analyzed using Seurat (5.1.0) package. Sample specific filtering criteria was applied to get the quality cells for clustering and annotation of scRNAseq data. Primary clusters of cells were identified based on following marker genes: Tumor cells (EPCAM, CD24), Stromal cells (PDPN, DCN), T cells (PTPRC, CD3D), Macrophages (PTPRC, CD14, CD68), Natural Killer cells (PTPRC, GNLY), Dendritic cells (PTPRC, CD1C), B cells (PTPRC, CD79A) and Plasma cells (MZB1). Differentially expressed genes were identified by ‘FindMarkers’ function of Seurat package using following criteria (logfc.threshold = 0.8, assay= “RNA”, min.pct = 0). Sub-clustering of T cells and macrophages were performed based on same steps as primary clustering and marker genes shown as dot plot in figure sections. No replicates were performed for the scRNAseq experiment.

### Isolation and identification of circulating tumor cells (CTCs)

The herringbone (HB) was designed using CAD software (AutoDesk AutoCAD) and manufactured using the photolithography method based on the previous report ^[35]^. A silicon wafer imprinted with the HB chip design served as the primary mold for fabricating PDMS structures. The PDMS gel was concocted by blending the elastomer base (Part A Sylgard 184) with an elastomer curing agent (Part B Sylgard 184) in a proportion of 10:1 (4019862, Dow, MI). Following plasma treatment, the HB-chips and PDMS base were conjoined to yield a complete set of PDMS HB-chips. These HB-chips were sequentially coated with poly (diallyl dimethylammonium chloride) (522376, Sigma Aldrich, MA) and a biotin-Alg solution (Bio-Alg) at concentrations of 2 mg/mL, resulting in layers bearing positive and negative charges. Subsequently, a biotin-avidin solution (PI31000, Molecular Probes, OR) at a concentration of 50 µg/ml was applied to the coated chips and incubated for a minimum of four hours. Capture antibodies were then introduced to these coated HB-chips. Biotin conjugated x-human anti-Doppel (A05457-Biotin-50ug, Boster Biological Technology, CA) and biotin conjugated x-human anti-EPCAM (51985, Cell Signaling, MA) antibodies were employed as capture antibodies within the HB-chips. Post antibody coating, the HB-chips were left to incubate overnight to facilitate effective binding between avidin and biotin. These HB-chips were then utilized to capture circulating tumor cells (CTCs) from patient blood samples. Whole blood samples were processed within 24 hours of collection from patients. These samples were prepared by incorporating 0.4 IU heparin (9041081, Thermo Fisher Scientific, MA) solution and diluting the blood with saline at a 1:1 ratio prior to introducing them to the capture antibody coated HB-chips. The prepared blood sample was channeled through the HB chips at a flow rate of 3 µl/min for a duration of 60 minutes, allowing a total of 200 µl of blood to traverse each HB chip. This procedure ensured minimal obstruction during the blood’s passage through the HB chips. For each blood sample, a minimum of five HB chips coated with Doppel and EPCAM were employed. Post-capture, the chips were rinsed (PBS with 0.1% BSA) at a flow rate of 10 µl/min to remove all non-captured cells. The captured cells were then fixed (4% formaldehyde) and permeabilized (goat serum and 0.3% Triton X-100) in preparation for staining. The cells were stained with DAPI (62247, Thermo Fisher Scientific, MA), cytokeratin (1:500, 502086852, Biolegend, CA), CD45 (1: 100, 502079010, Biolegend, CA) prior to imaging with an EVOS5000 fluorescence microscope. The CTCs were identified as DAPI+/CK+/CD45-.

### Statistical analysis

Where applicable, all experiments were conducted at least three times, and the mean and standard deviation (mean ± SD) were calculated. Statistically significant differences involving two groups were assessed using unpaired two-tailed Student’s t-tests. One way ANOVA was utilized to find statistical differences between more than two groups. Results were considered statistically significant at *p < 0.05, **p < 0.01, ***p < 0.001 and ****p < 0.0001 for all analyses.

## Data and code availability

All the raw FASTQ files, from both bulk and single-cell RNA sequencing study, supporting our hypothesis in this study will be made available in Gene Expression Omnibus (GEO) database upon accepting this article for publication. Customized ‘R’ script will be provided as supplemental file upon accepting this article for publication.

## Acknowledgements

This study was supported by grants from NIH R21CA264627, NIH R01CA262788, NIH SC1GM144171, DoD HT94252410217, DoD STTR Phase II #HT942523C0044, Lizanell and Colbert Coldwell Foundation #NAID20220187, TTUHSC-UTEP Seed Grant mechanism that are awarded to Dr. Taslim Al-Hilal. We obtained tissue microarray samples from US Biomax Inc. The authors like to thank Novogene Corporation Inc. for their support to perform RNAseq and scRNAseq.

## Author contributions

**TAA and FA** conceived the idea and planned the experimental designs. **ZA**, **XZ, RW, BK, FA, MMH and MH** performed most of the experiments. **XZ and RW** performed the ELISA experiment. **MMH** setup and prepared HB-µ chip for CTC capturing**. ZA, RW and MH** contribute to develop organoid model and processing the ascites for further experiment. **ZA** performed all bioinformatics analysis, RNAseq data analysis and scRNAseq data analysis. **JYZ** supplied the training cohort samples and helped in analyzing the samples. **BK and JYS** recruited the patients at Korea University Anam hospital, performed ELISA using first validation cohort samples and analyzed the data. **EPT and SYR** screened, recruited patients, and collected specimens at TTUHSC El Paso site. **ZA, MMH and TAA** wrote the manuscript. **ZA, JUC, MR, JYZ, SR, YB, ISK, and FA** discussed the data and contributed to writing the manuscript. **TAA** provided funding and supervised the entire study in conceptualization, designing, analyzing, writing. All authors contributed to manuscript revision and agreed on the final version of the manuscript.

