## Supplemental tables and figures for "The prion-like protein Doppel: A soluble biomarker steering ovarian cancer’s peritoneal to circulatory dissemination"

**Table S1:** Demographic information of recruited participants

| Demographics | Controls | Cases |
|---|---|---|
| N | **17** | **22** |
| Age, yr, mean (SD) | 50.4 (12) | 60.1 (14.1) |
| ≥ 55 years | 12 | 10 |
| < 55 years | 5 | 12 |
| Height, mean (SD) | 63 (2.2) | 67.9 (3.5) |
| Weight, mean (SD) | 179.4 (46.4) | 150.8 (23.5) |
| BMI, mean (SD) | 31.4 (6.2) | 26.5 (4.0) |
| **Ethnicity** | | |
| Hispanic | 16 (94.1%) | 18 (81.8%) |
| Non-Hispanic | 1 (5.8%) | 4 (18.2%) |
| **Race** | | |
| White | 15 (82.26%) | 19 (86.36%) |
| Other (mixed race) | 1 (5.88%) | 2 (9.1%) |
| Unknown | | 1 (4.55%) |
| Asian | 1 (5.88%) | |
| **Physical Activity** | | |
| Yes | 12 (70.6%) | 9 (40.9%) |
| No | 4 (23.5%) | 13 (59.1%) |
| Unknown | 1 (5.9%) | |
| **Family history of Ovarian cancer** | 0 (0.00%) | 1 (4.55%) |
| **Family history of Other cancer** | 9 (40.4%) | 6 (42.9%) |
| **FIGO Stage** | N/A | I (2), II (1), III (12), Benign (7) |

**Table S2**: Bio-banked ascites and Ascites-derived organoids (AsO) with clinical pathology

| | | | | Ascitic fluid | | | |
|---|---|---|---|---|---|---|---|
| **Pt. No.** | **Ethnicity** | **Race** | **EOC Type (FIGO Stage)** | **Doppel level** | **SC-SEQ** | **AsO established** | **RNA-Seq of AsO** |
| #27 | Hispanic | White | Mucinous Adenocarcinoma (IIB) | Low | Yes | Yes | Yes |
| #24 | Hispanic | Mixed race | Serous Grade 3 (IIIC) | High | Yes | Yes | Yes |
| #31 | Hispanic | White | Carcinosarcoma (IIIC) | High | N/A | N/A | |
| #40 | Hispanic | White | Low grade serous cancer (III A2) | High | N/A | N/A | |
| #44 | Hispanic | White | Mucinous Adenocarcinoma (IIA) | High | Yes | Yes | Yes |
| #46 | Hispanic | White | Serous Grade 3 (IIIA) | No Ascites | | | |
| #47 | Hispanic | While | Serous Grade 3 (IV) | No Ascites | | | |
| #48 | Hispanic | White | Serous Grade 3 (IIIC) | Low | Yes | Yes | Yes |

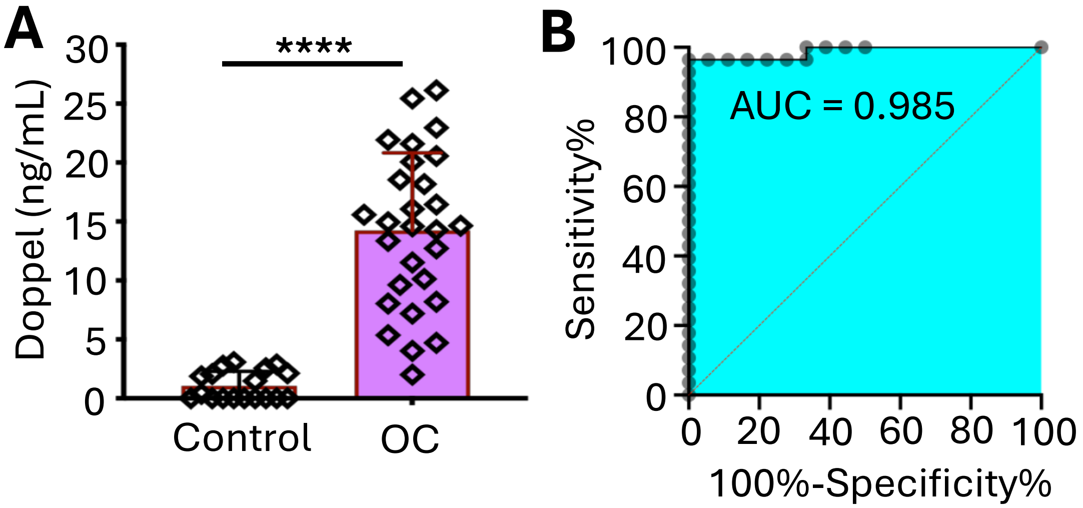


**C**

| | Control | OC |
|---|---|---|
| **Cases No.** | 18 | 28 |
| **Mean (ng/mL)** | 1.06 | 14.24 |
| **SD** | 1.23 | 6.58 |
| **Positive** | 0/18 | 26/28 |

**Figure S1: Serum Doppel level is elevated in training set of ovarian cancer patients. A)** Measurement of serum Doppel of ovarian cancer and control subjects by ELISA. Error bar represents mean ± SD. **B)** The ROC curve of Doppel between control and ovarian cancer serum samples. **C)** Serum Doppel level comparison chart between control and ovarian cancer subjects. **** $p$= <0.0001.

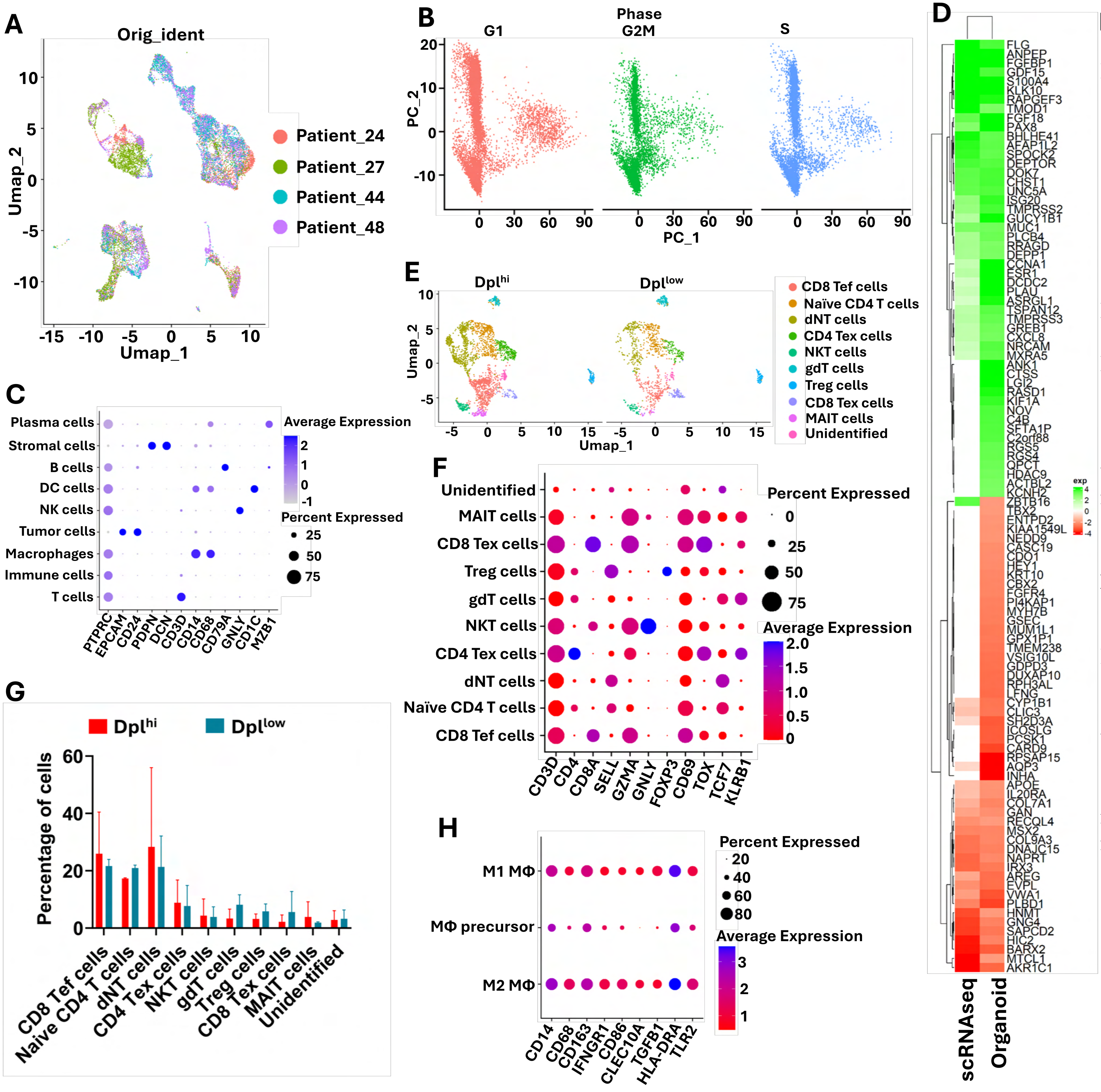


### Figure S2

**Figure S2: Single-cell RNA sequencing analysis of Doppel high and low ascites derived cells group. A)** UMAP plot showing the effective uniform distribution of cells after integration of four individual scRNAseq datasets. **B)** Cell cycle distribution of four integrated scRNAseq datasets. **C)** Heat map comparison of top 50 up- and down-regulated genes in Doppel high group from organoid with genes differentially expressed genes in Doppel high tumor cells subcluster from scRNAseq data. Log2 fold changes are included in this analysis. Genes not found in scRNAseq analysis shown as 0 value in this heat map. **D)** Dot plot showing the marker genes expression in each identified primary cluster. **E)** UMAP plot showing the distribution of T cell populations between Doppel high and low ascites derived cell group. Secondary analysis was performed on T cells cluster. **F)** Dot plot showing the marker genes expression in each identified secondary T cells sub-cluster. **G)** Bar chart showing the percentage of cells, in secondary T cells clusters, present in each ascites derived cell group. Error bar represents mean +SD. **H)** Dot plot showing the marker genes expression in each identified secondary macrophages sub-cluster.
